# Systemic immune dysregulation in hypertensive disorders of pregnancy persists years after delivery

**DOI:** 10.1101/2025.11.25.690209

**Authors:** Maximilian Sabayev, Edward A. Ganio, Ina A. Stelzer, Masaki Sato, Amélie Cambriel, Thomas A. Bonham, Zala N. Juvan, Maïgane Diop, Purnima Iyer, Oshra Sedan, Nima Aghaeepour, Martin S. Angst, Gary M. Shaw, David K. Stevenson, Marcia L. Stefanick, Heather A. Boyd, Mads Melbye, Mark A. Hlatky, Virginia D. Winn, Brice Gaudilliere, Dorien Feyaerts

## Abstract

**Background:** Hypertensive disorders of pregnancy (HDP), including preeclampsia and gestational hypertension, are associated with an increased risk of cardiovascular disease (CVD) later in life. Mechanisms that link HDP to CVD, however, remain unclear.

**Methods:** We used a high-dimensional single-cell mass cytometry approach to profile the distribution and functional responses of maternal immune cells in three separate groups of HDP cases and normotensive controls, sampled antepartum, postpartum, and several years postpartum (midlife). We used multivariable sparse modeling to distinguish HDP cases from controls.

**Results:** We accurately distinguished HDP cases from controls at all three study timepoints, with area under the receiver operator characteristic (AUROC) curve values of 0.814 for the antepartum group, 0.757 for the postpartum group, and 0.692 for the midlife group. Distinct immune signatures for each model underscore the dynamic dysregulation of the immune system throughout life. In addition, we identified a persistent immune dysregulation signal among HDP cases at all three timepoints, characterized by increased B cell frequency and decreased pSTAT3 response upon cytokine stimulation in classical monocytes.

**Conclusions:** Persistent immune dysregulation among women with a history of an HDP may contribute to elevated long-term risk of CVD development.

## INTRODUCTION

Hypertensive disorders of pregnancy (HDP), which include preeclampsia and gestational hypertension, affect approximately 5% to 8% of pregnancies and are a leading cause of maternal and perinatal morbidity and mortality worldwide.(1) While the clinical manifestations of HDP typically resolve shortly after delivery, women with a history of HDP face significantly increased risks of developing cardiovascular disease (CVD) and major adverse cardiovascular events, such as myocardial infarction and stroke, later in life (1–3).

Despite these epidemiological links, the mechanisms by which HDP predisposes women to the subsequent development of CVD remain unclear. Although several biological factors likely play a role, recent evidence suggests that persistent immune system alterations may be a key driver of CVD in patients who have had HDP (4). Preeclampsia and gestational hypertension are associated with profound innate and adaptive immune cell dysfunction (5–7). At the maternal-fetal interface, decreased uterine natural killer (NK) cell numbers (8), altered NK cell receptor expression (8), activated and pro-inflammatory macrophages (8, 9), and reduced numbers of regulatory T cells (Treg) (10, 11) have been reported in patients with preeclampsia, and may contribute to inadequate spiral artery remodeling and impaired immune tolerance towards the fetus in approximately 15% of cases. Preeclampsia is also associated with systemic immune dysregulation, including decreased peripheral Treg frequencies (6, 12), reduced CD4 T cell function (5), and increased pro-inflammatory monocytes (9). Interestingly, systemic immune dysregulation has been observed after the delivery of the fetus, well into the postpartum period, including increased inflammatory mediators (13, 14) and reduced proportions of activated CD4^+^ memory T cells (15), raising the hypothesis that persistent immune dysregulation after pregnancy contributes to the increased risk of developing CVD.

The Effect of Preeclampsia on Cardiovascular Health (EPOCH) study was designed to address this hypothesis by longitudinally investigating biological and immune profiles in women with and without a history of HDP (2, 16–18). By assessing whether certain biological determinants are persistently altered in women with HDP, the EPOCH study aims to determine the relationship between specific biological alterations observed during this event and the potential predisposition to CVD years later.

Mass cytometry is a powerful method to comprehensively characterize human immune responses (19–21) at the single-cell level, and has been used to study both healthy and complicated pregnancies (5, 22, 23). Mass cytometry allows for the simultaneous measurement of over 40 proteomic parameters per cell analyzed, enabling the precise phenotyping of all major immune cell populations present in a blood sample as well as their functional state, represented by the phosphorylation of intracellular signaling proteins (19, 20). Mass cytometry has been utilized to decode the maternal immune clock of pregnancy (22, 23) and identify immune signaling responses within innate and adaptive immune cells that can predict the development of preeclampsia before the onset of clinical symptoms (5).

In this study, we utilized a high-dimensional, single-cell mass cytometry approach to monitor the distribution and function of major peripheral immune cell subsets (i.e. the maternal immunome) across three timepoints in women with and without an HDP: during pregnancy (antepartum), at least six weeks after delivery (postpartum), and more than two years after delivery (midlife). Combined with multivariable sparse modeling, this approach allowed us to identify immunome changes over the years immediately following delivery and to detect persistent changes that may mediate the observed link between HDP and future CVD risk.

## METHODS

### Study design

The Effect of Preeclampsia on Cardiovascular Health (EPOCH) study, which is investigating the long-term effects of preeclampsia on women’s cardiovascular health outcomes, has been previously described (2, 16–18). This study was approved by the Stanford University Institutional Review Board, and all participants provided written informed consent.

We recruited two groups of women: one during pregnancy, and the other at least two years after delivery. In the pregnancy group, we screened women aged 18 to 45 years receiving prenatal care at Stanford University Medical Center for cases with new-onset HDP and recruited normotensive pregnant women matched for maternal age and gestational age as controls. We excluded participants with chronic hypertension, diabetes, heart disease, chronic kidney disease, autoimmune disease, or cancer before pregnancy, and excluded pregnancies with known fetal chromosomal or structural anomalies. Furthermore, controls were excluded if they developed preeclampsia, gestational hypertension, or gestational diabetes after study recruitment. We defined preeclampsia and gestational hypertension using the 2013 American College of Obstetricians and Gynecologists criteria (24). Participants in the pregnancy cohort were invited to complete two study visits, the first antepartum and the second at least six weeks postpartum. However, due to COVID-19 pandemic restrictions, only 17 cases and 33 controls were able to return for their second study visit, while 25 cases and 4 controls were only able to participate in the postpartum visit and did not attend the antepartum visit.

We recruited a separate group of women who were at least two years postpartum (midlife group) by searching the electronic health records of the Stanford Health Care system for pre-menopausal women between 21 and 55 years of age who had delivered a baby at Stanford at least two years earlier and met the inclusion and exclusion criteria described above. Case women had their most recent pregnancy complicated by preeclampsia and control women were matched to the cases on maternal age and time since delivery.

### Single-cell analysis with mass cytometry

#### Ex vivo immunoassay

We drew fasting whole blood samples into a sodium-heparinized vacutainer at each study visit. We created four aliquots of whole blood and stimulated them for 15 minutes at 37°C with either lipopolysaccharide (LPS) (1μg/mL), interleukin (IL)-18 (100ng/mL), a cocktail containing IL-2, IL-4, and IL-6 (100ng/mL each), or phosphate-buffered saline (PBS; unstimulated condition) to activate canonical pregnancy related signaling responses ex vivo (5, 22, 23). We then fixed the samples with Proteomic Stabilizer (SmartTube) for 10 minutes and stored them at -80°C until further processing.

#### Derivation of cell frequency and functional response features

After thawing samples, we lysed red blood cells and then processed them for barcoding, staining, and analysis, as previously described (25, 26). We acquired data from mass cytometry by time-of-flight (CyTOF) using the Helios^TM^ CyTOF system (Standard BioTools). We applied a 45-marker antibody panel, which included 26 surface and 19 intracellular markers (**Table S1**), to measure the frequency of innate and adaptive immune cells, as well as the signal intensity of phosphosignaling proteins and functional markers. We identified innate and adaptive immune cells using a manual gating strategy, as shown in **Figure S1**.

We report cell frequencies as a percentage of mononuclear cells, with the exception of granulocyte subsets, which we report as a percentage of singlet leukocytes. We measured signal intensities of phosphosignaling proteins and functional markers using the arcsinh transformed value (arcsinh(x/5)) from the median signal intensity (19). We assessed the endogenous signaling/functional activity by analyzing unstimulated cells and expressed signaling/functional responses to stimulation as the arcsinh transformed ratio relative to the endogenous signaling/functional response, i.e. the difference in arcsinh transformed signal intensity between the stimulated and unstimulated condition. We applied a penalization matrix to the dataset (**Suppl. File SF1**) based on prior mechanistic immunological knowledge of receptor-specific canonical signaling pathways (27). In this matrix, a value of 1 indicates features that were included in the analysis, while a value of 0 indicates those that were excluded. As such, the penalization matrix emphasizes the combination of immune cells and their signaling/functional responses used in the subsequent analyses. This approach resulted in 2,172 features for each sample: the frequencies of 35 innate and adaptive immune cells, 700 endogenous response features, 248 LPS-induced response features, 635 IL-2/IL-4/IL-6-induced response features, and 554 IL-18-induced response features.

#### Data visualization

We visualized the complete single-cell immune dataset, comprising all three groups of participants, using uniform manifold approximation and projection (UMAP) embedding. We indicated cell populations on UMAP using the labels from the manual gating approach. We used a *t*-distributed stochastic neighbor embeddings (t-SNE) layout to visualize the measured features in each group; as such, each feature is represented in triplicate, once for each group.

### Statistical analysis

#### Multivariate modeling

We employed a sparsity-inducing multivariable modeling approach to analyze the high dimensional mass cytometry data and account for the large number of immune features relative to the number of samples included in the study. We fitted sparse classification models independently to each group to discriminate between HDP cases and controls. We applied several logistic regression-based classifier candidate models to each dataset, including least absolute shrinkage and selection operator (LASSO), adaptive LASSO, and their Stabl-enhanced counterparts (28). Additionally, we integrated the immune feature datasets using either a late fusion or early fusion approach (28). In the late fusion approach, we first segregated the immune feature dataset based on stimulation condition (frequency, unstimulated, LPS-stimulated, IL-2/IL-4/IL-6-stimulated, and IL-18-stimulated response), fitted models for each stimulation dataset separately, and then combined the results into a final model. In the early fusion approach, we included all immune features into a single model. We assessed predictive performance using cross-validation through a 200-times repeated 5-fold cross-validation approach, in which at each fold 80% of sampled individuals were randomly designated as the training set and the remaining 20% individuals were used for validation testing. Using the predictions on the test data within cross-validation, we calculated area under the receiver operator characteristic (AUROC) curve and assessed statistical significance by Mann-Whitney U-test p-values, considering a p-value < 0.05 to be statistically significant. In the manuscript, we report the model with the highest AUROC (**Suppl. File SF2**). The informative features selected by the model were determined by fitting the model a final time on the whole dataset.

#### Confounder analysis

We performed a post-hoc confounder analysis (**Table S2**) to assess the potential confounding effect of body mass index at study visit, parity, maternal age, gestational diabetes, blood pressure, and timing of study visits on the predictive accuracy of the models. This analysis involved fitting a logistic regression model that incorporated both the potential confounding variables and the model predictions generated from the cross-validation. We calculated F-statistics on the coefficients of each confounding variable to test whether it accounted for the model predictions.

#### Temporal analysis

After being trained on a given timepoint, each of the multivariate models were used to discriminate between cases and controls at chronologically later timepoints. The model trained on the antepartum group was applied on the postpartum and midlife groups, and the model trained on the postpartum group was applied on the midlife group.

#### Identification of overlapping features across groups

We used a univariate approach to identify persistent immune features that were differently expressed across more than one cohort. We applied the Mann-Whitney U-test to identify significantly different features between cases and controls for each cohort separately.

### Data availability

Raw (FCS files), processed data (cell frequency and phosphosignaling/functional response), and patient characteristics will be made available on Dryad after peer review and upon publication. Any additional information required to reanalyze the reported study is available upon request from the corresponding author.

## RESULTS

### Study population and study design

We studied three distinct groups of participants recruited between June 2019 and November 2023: antepartum, postpartum, and midlife (**Figure 1**). Cases were more likely than controls to have a higher body mass index (BMI), gestational diabetes, and higher blood pressure at all three study timepoints (**Table 1)**. In the postpartum group, the first study visit occurred earlier for controls than cases. There were no differences in maternal age at study visit, parity, or the proportions of nulliparous and primiparous women among the three study groups.

**Figure 1.**
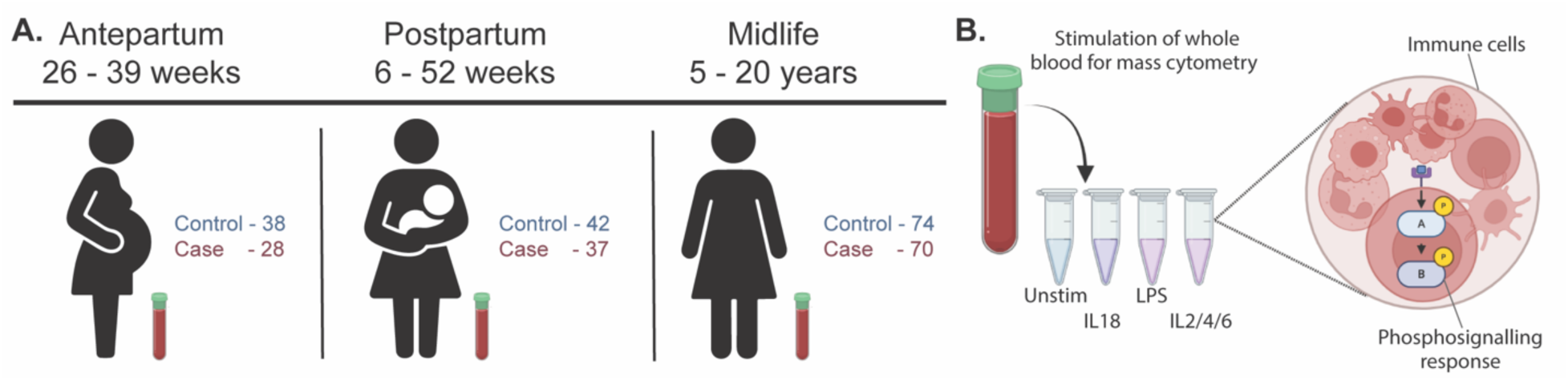
Study overview. **A.** Three distinct groups were studied. The antepartum group was enrolled between 26 and 39 weeks of gestation. The postpartum group was enrolled 6-52 weeks after delivery, and the midlife group 2-20 years since last delivery. **B.** Peripheral blood samples were collected from each participant for the assessment of major immune cell frequencies and functional responses using single-cell mass cytometry. Blood samples were either left unstimulated or stimulated with interleukin (IL)-18, lipopolysaccharide (LPS), or a cocktail of IL-2, IL-4, and IL-6.

**Table 1.**
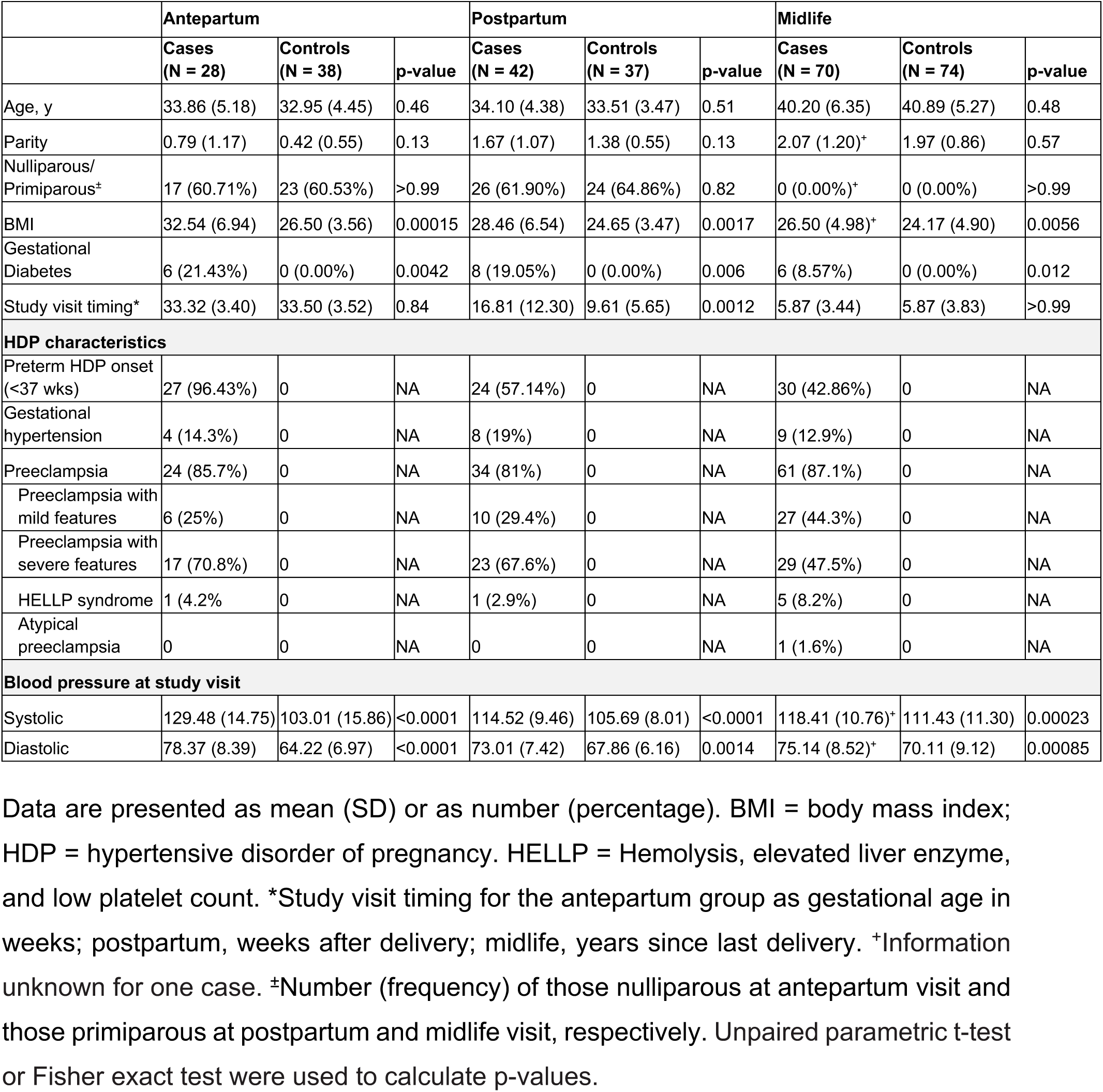
Demographics and clinical information of cases and controls per group at study visit.

The major peripheral innate and adaptive immune cells (i.e. the maternal immunome), assessed by mass cytometry, from all three groups of participants are displayed in a UMAP layout to illustrate their distribution and cell- and stimulation-specific phosphosignaling responses (**Figure 2**). The UMAP representation confirms the canonical stimulation responses in study participants, with examples including increased phosphorylated Cyclic AMP Response Element Binding Protein (pCREB) expression in monocyte subsets after LPS stimulation, increased pCREB expression in natural killer (NK) and NKT-like cells after IL-18 stimulation, and phosphorylated signal transducer and activator of transcription 3 (pSTAT3) expression in monocytes and T cell subsets after IL-2/IL-4/IL-6 stimulation.

**Figure 2:**
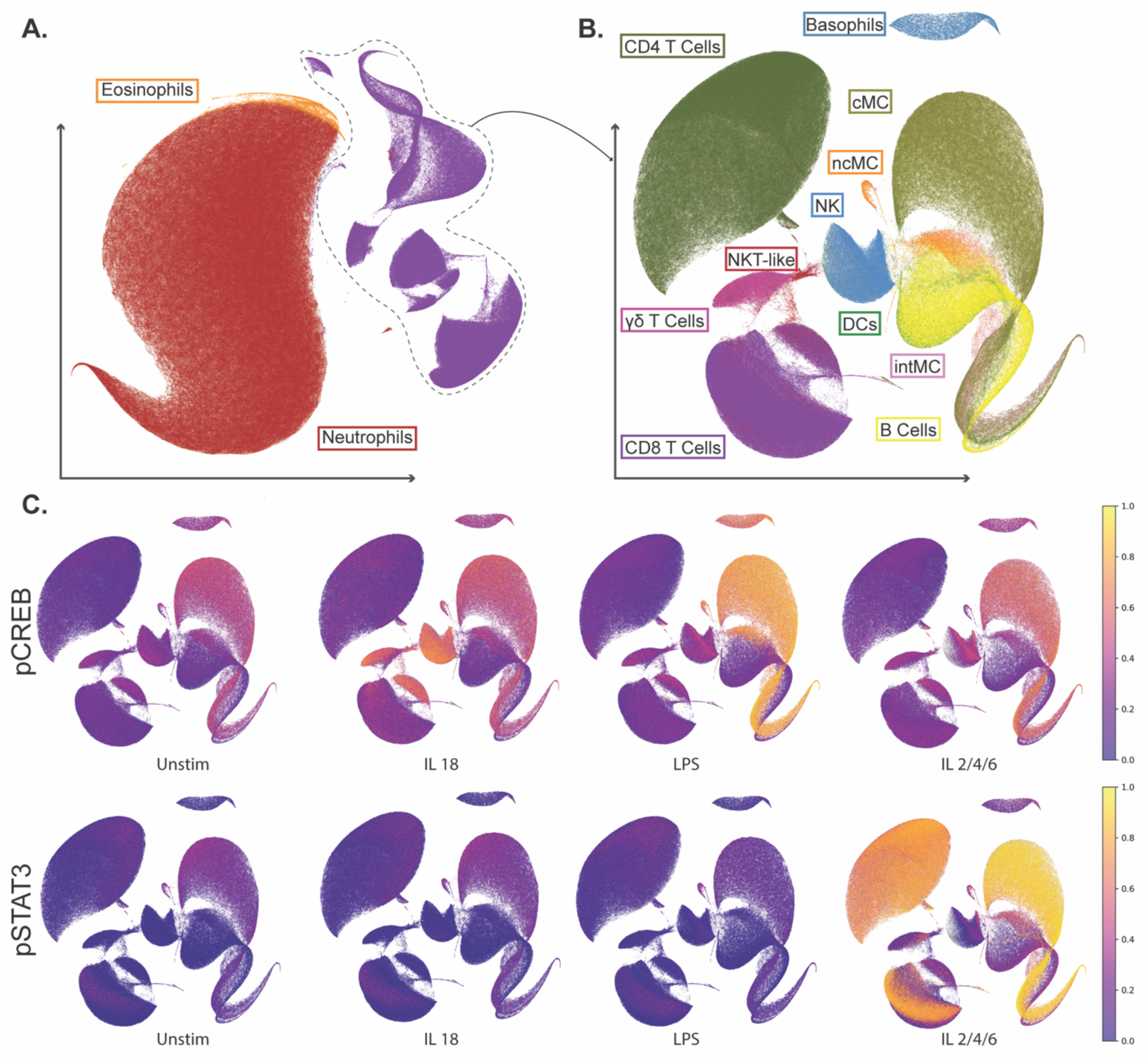
Single-cell mass cytometry analysis of the maternal immunome highlights cell type- and stimulation-specific phosphosignaling responses. **A-B.** Uniform manifold approximation and projection (UMAP) representation of the single-cell mass cytometry dataset of all three groups combined. UMAP is a dimensionality reduction technique that allows us to generate two-dimensional embeddings for cells with similar marker expression together, thereby allowing the identification of cell clusters within the data visually. **A.** UMAP represents all live leukocytes, including eosinophils, neutrophils, and mononuclear cells. **B.** UMAP representation of only mononuclear cells. Gated cell populations are annotated. **C.** UMAPs representing mononuclear cells are colored according to intracellular phosphosignaling response at baseline (Unstim), and after in vitro stimulation with IL-18, LPS, and a cocktail of IL-2/IL-4/IL-6. Z-scored median expression of phosphorylated CREB (pCREB) and phosphorylated STAT3 (pSTAT3) are shown as examples. cMC = classical monocyte. ncMC = non-classical monocyte. intMC = intermediate monocyte. DCs = dendritic cells. NK = natural killer cell. NKT-like = natural killer T cell-like.

### Distinct immune signatures identify hypertensive disorders of pregnancy across three timepoints

A manual gating strategy identified 35 immune cell subsets and their functional responses, resulting in a final dataset comprising 2,172 immune features per sample. Sparse multivariable predictive modeling discriminated HDP cases from controls with high predictive accuracy at each study timepoint (**Figure 3)**. In the antepartum group, the AUROC was 0.814 (p-value = 0.00002); in the postpartum group, the AUROC was 0.757 (p-value = 0.00009); and in the midlife group, the AUROC was 0.692 (p-value = 0.00007) (**Suppl File SF2**). A confounder analysis demonstrated that all three models remained significantly predictive of HDP when potential confounders such as BMI at study visit, parity, maternal age, gestational diabetes, blood pressure, and timing of study visits were accounted for (**Table S2**).

**Figure 3:**
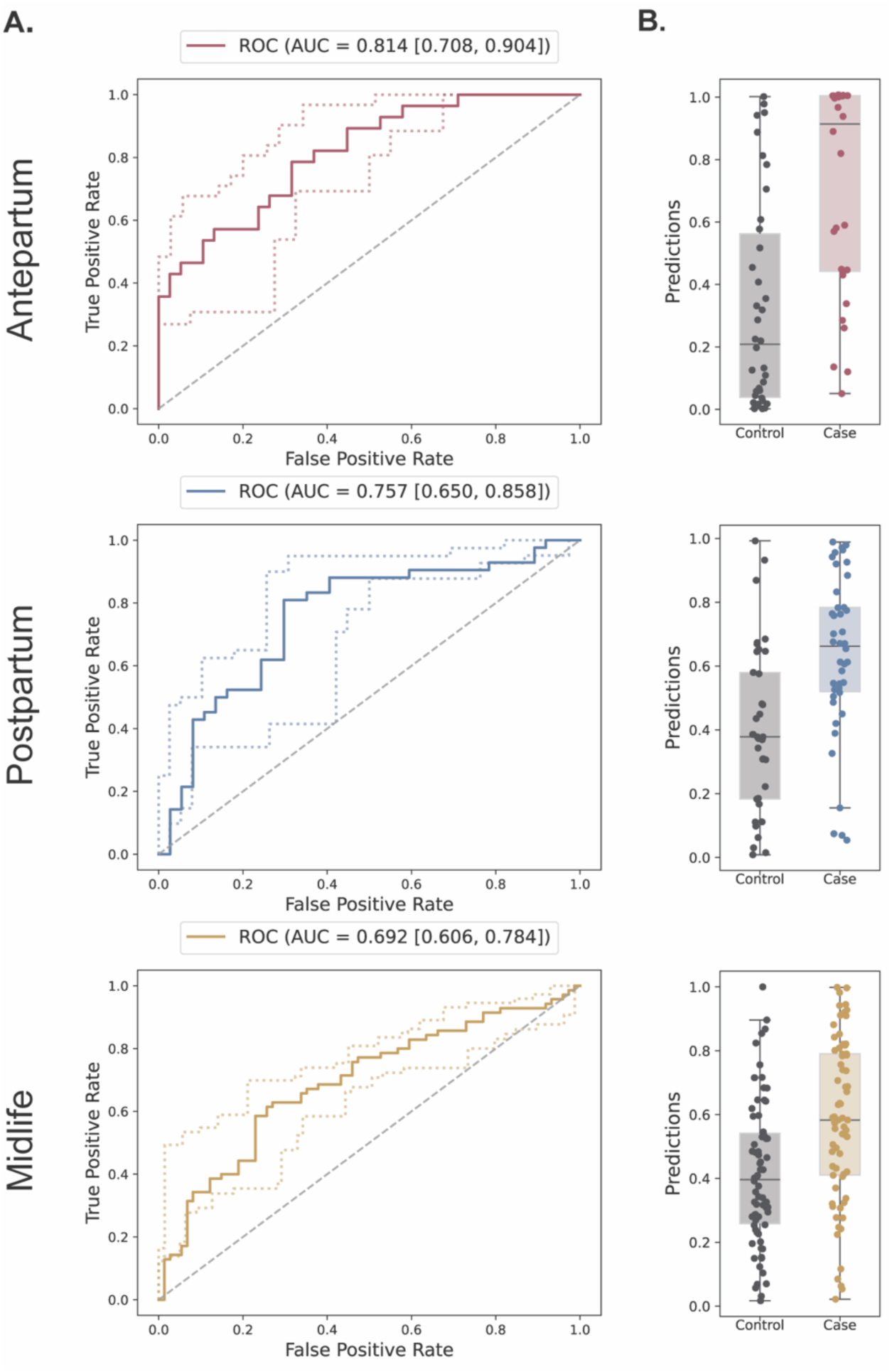
Multivariable sparse modeling of immune profiles classifying HDP cases and controls across the lifespan. The results for each group – antepartum, postpartum, and midlife – are shown in their respective rows. **A.** Receiver Operator Characteristics (ROC) of the cross-validated predictions of the best performing models in each group. The Area Under the Curve (AUC) is calculated along with its 5% confidence interval quartiles. **B.** Boxplots of the cross-validated predictions in each group.

A distinct immune signature of HDP was evident at each study timepoint, with 37 features selected in the antepartum model, four features in the postpartum model, and 13 features in the midlife model (**Table S3**), showing no overlap of specific features among the three models. Seventeen features (46%) from the antepartum model pointed to altered metabolic and functional responses to IL-18 stimulation in various innate and adaptive cells. For example, IL-18 stimulation resulted in an increased glycolytic response in cases compared to controls, as evidenced by increased glucose transporter 1 (GLUT1) and hexokinase 2 (HK2) expression. Furthermore, markers of cell chemotactic capacity, including CD62L and CXCR4, were decreased in cases in both the antepartum and postpartum groups (**Suppl Fig S2A-B**). In addition, cases in the midlife cohort showed a decreased pCREB and pSTAT6 response to both LPS and IL-2/IL-4/IL-6 stimulation in innate immune cells, decreased CXCR4 expression by Ki67+ CD8T+ cells upon IL-2/IL-4/IL-6 stimulation, and an increased expression of the fatty acid oxidation marker carnitine palmitoyltransferase 1A (CPT1A) in regulatory T cells (**Suppl Fig S2C**). This disconnect in specific immune features was further supported by the observation that while the antepartum model was able to discriminate HDP history in the postpartum cohort (AP➔PP: AUROC = 0.713, p-value = 1.17e-6), neither the antepartum nor postpartum model could predict HDP history for the midlife cohort (AP➔ML: AUROC = 0.527, p-value = 0.577; PP➔ML: AUROC = 0.571, p-value = 0.144) (**Suppl. Fig. S3**). This indicates that while the immune signature identified at the time of acute HDP remains predictive into the postpartum period, it does not extend its predictability into midlife. These findings reveal timepoint-specific immune signatures that reflect the evolving dysregulation associated with HDP during pregnancy, postpartum, and even years later.

### Persistent immune dysregulation associated with hypertensive disorders of pregnancy

While the multivariable approach offered a statistically stringent comparison between cases and controls, the sparse feature selection may have excluded differentially expressed immune markers that were either not major contributors to the overall models or were correlated with the selected features. To explore persistent immune dysregulation across all time points more comprehensively, we applied a less stringent univariate statistical approach. Immune features differentially expressed between cases and controls were represented on a Venn diagram, highlighting shared features between the three timepoints (**Figure 4**). While the majority of differentially expressed immune features were specific to the individual group, 75 immune features (17%) were differentially expressed between cases and controls at several study timepoints (**Figure 4A, Suppl File S2**). Between the antepartum and postpartum group, 53 features were shared, with 21 features (40%) indicating a decreased expression of CXCR4 upon IL-18 stimulation in T cell and innate cell subsets in cases (**Figure 4B**). Ten features were shared between the postpartum and midlife group, four of which suggested that a history of HDP was associated with decreased pCREB signaling in response to LPS stimulation in innate immune cells. Sixteen features were shared between the antepartum and midlife groups, seven of which indicated increased endogenous pSTAT3 activity in T cell subsets. Interestingly, two immune features were differentially expressed across all three groups: the frequency of B cells was consistently increased in cases, whereas pSTAT3 expression after IL-2/IL-4/IL-6 stimulation in classical monocytes was consistently decreased in cases. These findings emphasize the dynamic nature of the immunome in women with HPD and reveal cell-specific and persistent dysregulation of the systemic immune system that might contribute to the increased risk of CVD.

**Figure 4:**
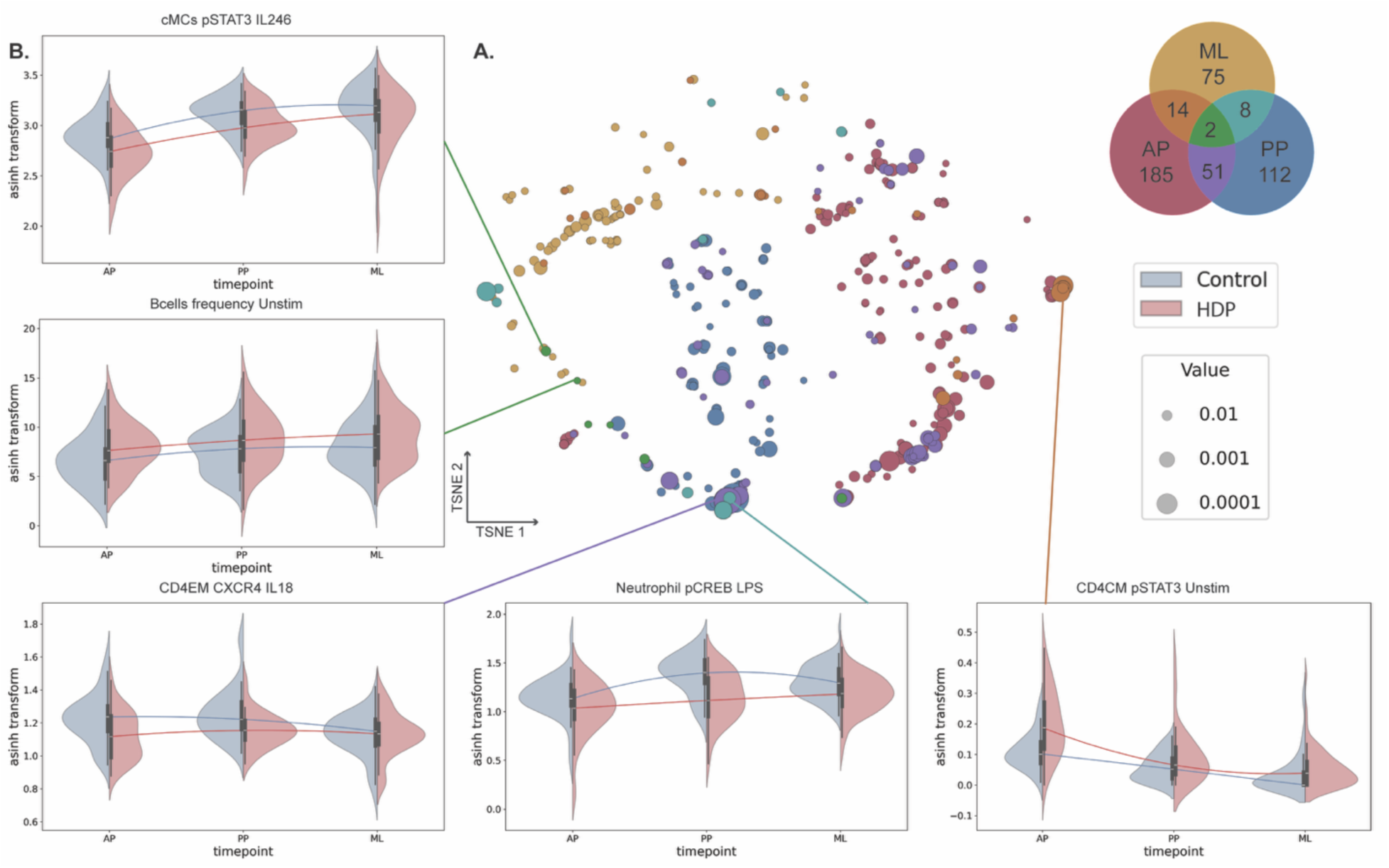
Univariate analysis reveals a persistent immune signature of HDP. **A.** Immune features differentially expressed between cases and controls are represented on a t-SNE embedding and were determined via Mann-Whitney U-tests within each group, with significance being p-values < 0.05. Node size reflects the related Mann-Whitney U-test p-value. Nodes are colored based on the colors represented in the Venn diagram, which depicts the number of differently expressed features that are shared across the three groups. **B.** Violin plots highlight representative features, either median marker expression or frequency. The interpolated median expression across the groups is highlighted by the line in a darker shade. AP = antepartum; PP = postpartum; ML = midlife.

## DISCUSSION

In this study we conducted an extensive single-cell analysis of the maternal peripheral immune system to identify evolving and persistent immune signatures associated with having an HDP. Sparse multivariable modeling showed that the maternal immune system evolves over time in women with a history of HDP compared with normotensive controls, exhibiting distinct differences not only during pregnancy, but also throughout the postpartum period and extending several years thereafter. Moreover, differentially expressed immune features indicated that cell-type specific immune dysregulation observed during a pregnancy complicated by an HDP may persist long after pregnancy, and could contribute to the development of CVD later in life.

The multivariable analysis of the immunome dataset highlighted that the maternal peripheral immune system can accurately differentiate HDP from normotensive pregnancies during, shortly after, and several years after pregnancy. While there was no overlap of specific features selected among the three models, two general themes consistently emerged in the HDP-associated immune dysregulation at each time point: an altered immunometabolism and diminished immune cell chemotactic capability. In samples taken during pregnancy, IL-18 stimulation increased expression of GLUT1 and HK2 in innate and adaptive immune cells in HDP cases compared with controls. These elevations enable immune cells to rapidly enhance their glycolytic capacity, a sign of immune cell activation and heightened inflammatory responses (29). Increased endogenous expression of CPT1a by regulatory T cells in HDP cases at midlife and during pregnancy suggest an upregulation of fatty acid oxidation for energy production (30). Overexpression of CPT1a can lead to increased production of reactive oxygen species as a byproduct (31), thereby contributing to oxidative stress, which can be a risk factor for the development of CVD (32, 33). In addition, model features for all three groups showed a decrease in the expression of chemotactic markers CXCR4 and CD62L upon cytokine stimulation of various immune cells in HDP cases, in concordance with others (34). CXCR4 and CD62L expression are essential for immune cells to migrate to sites of inflammation (35–37). Reduced expression could impact their migration capability and subsequent resolution of inflammation. Moreover, CXCR4 has been shown to control tissue regeneration (36). As such, a reduced capacity of immune cells to upregulate CXCR4 in response to cytokines could indicate a diminished ability to repair vascular injury during inflammation in HDP cases.

Interestingly, all features in the postpartum model, as well as 46% of the features in the antepartum model, were part of the IL-18 stimulation dataset. Elevated serum and placental levels of IL-18 have been observed in preeclampsia cases (38) and IL-18 has been shown to induce endothelial damage (39). The altered response of the immune system towards IL-18 in women who had an HDP may indicate a central role for IL-18-mediated disruption of immune regulation and the overall pathophysiology of HDP.

Complementing the multivariate modeling, univariate analyses highlighted persistent and cell-specific immune dysregulations in cases compared with controls. Notably two immune features differed between the HDP and control groups at all time points: B cell frequencies and the pSTAT3 response in classical monocytes. B cells have emerged as important immune contributors to the development of CVD. More specifically, B1 cell subsets and regulatory B cells have been shown to play a protective role through the secretion of anti-inflammatory cytokines, while B2 cell subsets and plasmablasts enhance atherosclerosis development by promoting inflammation (40). Therefore, a persistently increased presence of B cells over time in women with a history of HDP may produce a long-term inflammatory state that promotes atherosclerosis and subsequent CVD. The idea of an immunoregulatory imbalance is further supported by Salby et al.’s findings of a reduced expression of immunosuppressive PD-1 on total B cells and regulatory B cells (Breg) in women with preeclampsia (41). In addition, STAT3 signaling plays a central role in the pathophysiology of vascular diseases, promoting inflammation in part by activating pro-inflammatory monocytes (42, 43). Paradoxically, we observed a decreased pSTAT3 response to inflammatory cytokine stimulation in patients with HDP. However, the observed dampened pSTAT3 response to ex vivo stimulation may be attributed to an elevated baseline pSTAT3 state caused by a heightened endogenous inflammatory cytokine milieu, which results in a diminished capacity to respond to additional ex vivo stimulation.

Multiple studies have examined the antepartum immune system at the onset of preeclampsia, comparing it to gestational age-matched controls using cytometry methods. Notably, only one study employed mass cytometry instead of flow cytometry, allowing for a more comprehensive analysis. The flow cytometry studies, while informative, are somewhat limited in their exploration of the broader immune system. They primarily focused on differences in a select set of immune cell frequencies, particularly within the Th17/Treg and Th1/Th2 paradigms (44). These paradigm findings support a shift towards decreased immunosuppression and increased pro-inflammatory responses in women with preeclampsia. Specifically, increased frequencies of Th17 cells (CD4+IL17A^+^ or RORγT^+^CD4^+^ T cells) (45–48) have been observed at the onset of preeclampsia, alongside decreased Treg frequencies (CD4^+^CD25^+^CD127^low^, CD4^+^FoxP3^+^, or CD4^+^CD25^+^FoxP3^+^) (10, 12, 45–56). Further evidence for this shift can be found in increased frequencies of Th1 cells (47, 57), exhausted Tregs (51), M1-like (9) and M2-like monocytes (58), and decreased frequencies of myeloid-derived suppressor cells (59), as well as tolerance-promoting Galectin-1^+^ and TIM3^+^ T cells and NK cells in women with preeclampsia (56, 60). In a unique mass cytometry study (5), the dynamic nature of the systemic immune system was characterized by collecting peripheral blood mononuclear cells during the first and second trimesters, prior to the onset of preeclampsia. This study demonstrated that dynamic changes in peripheral immune cell signaling responses during pregnancy, particularly impaired pSTAT5 signaling in CD4^+^ T cells, can differentiate women who later developed preeclampsia from controls before the onset of clinical symptoms, achieving an area under the curve of 0.91.

Few studies have examined immune cell differences in the postpartum period. One two-color flow cytometry study showed a decreased frequency of CD25^+^ T cells one week post-delivery in preeclampsia cases compared to normal pregnancies (61). In addition, at least six months postpartum, lower proportions of activated CD4^+^ memory T cell subsets were observed in women who had preeclampsia compared to controls (15). Our study differs from these earlier findings by including three different timepoints, examining the immune system more broadly, and emphasizing functional read-outs.

This study has certain limitations. Firstly, due to COVID-19 pandemic restrictions, patients included in the three groups only partially overlapped, precluding the longitudinal analysis of an individual’s immune responses over time. Secondly, while we found differentiating immune signatures during pregnancy, postpartum, and at midlife, we cannot conclude that these immune signatures originated during pregnancy. Future studies that include samples collected before pregnancy would be important to examine the contribution of pre-pregnancy immune dysregulation. Thirdly, we did not have enough women with HDP to examine specific differences among HDP-subtypes, such as preeclampsia versus gestational hypertension, early versus late onset, and preeclampsia with severe features versus preeclampsia without severe features. Moreover, while the mass cytometry immunoassay panel allowed for high resolution phenotypic and functional analysis of all major immune cell subsets, certain immune cell subpopulations were under-characterized. In particular, a more in-depth analysis of B cell subsets would enhance the identification of B cell-specific biomarkers associated with the development of CVD (40). Lastly, clinical assessment of CVD could not be performed, as none of the midlife participants, who were relatively young (mean age 41 years) (3, 62), had developed CVD at the end of the study. Future studies that include longer follow-up periods later in life should cover a higher incidence of CVD.

In summary, we found that the maternal immunome differs significantly between women with an HDP and normotensive controls, not only during pregnancy but also in the postpartum period and several years thereafter. The specific immune features vary by study timepoint, underscoring the dynamic response of the immune system throughout a woman’s life. Moreover, we documented the persistent immune dysregulation of two immune features in women with a history of an HDP, which may elucidate the mechanisms that connect HDP to the heightened risk of CVD later in life. These results highlight the need for more in-depth immune assessments and ongoing monitoring to identify targeted interventions that could help mitigate long-term health risks associated with HDP.

## Supporting information

Penalization matrix

Cross validation results

Univariate results

## ACKNOWLEDGEMENTS

We would like to thank Ijeoma Iwekaogwu, Ana A. Campos, Nick Bondy, Sonia Chavez, Elizabeth Sherwin, Christine Lee, Mira Diwan, and Maria Navarro for their efforts in enrolling participants and coordinating study visits; Archana Bhat, Katherine Connors, and John Maul for their assistance in identifying and contacting potential midlife study participants; Emma Anushka DuBridge and Adrian G. Yabut for processing samples; and Cele C. Quaintance for their support with study operations. We used Stanford Health Care and Stanford School of Medicine Secure GPT (beta, 2025) for language editing assistance. Finally, we are extremely grateful to the participants, whose involvement made this study possible.

## SUPPLEMENTAL MATERIAL

Figures S1-S3

Tables S1-S3

Files S1-S3

## NON-STANDARD ABBREVIATIONS AND ACRONYMS

AUROC: area under the receiver operator characteristic
AP: antepartum
EPOCH: Effect of Preeclampsia on Cardiovascular Health
ML: midlife
PBS: phosphate-buffered saline
PP: postpartum

