## Supplemental figures and tables for "Systemic immune dysregulation in hypertensive disorders of pregnancy persists years after delivery"

*Sabayev et al.*

Figure S1 - Immune cell gating strategy
Figure S2 - Informative model features
Figure S3 - Model performance when applied to later timepoints

Table S1 - Antibody panel
Table S2 - Confounder analysis
Table S3 - Informative model features

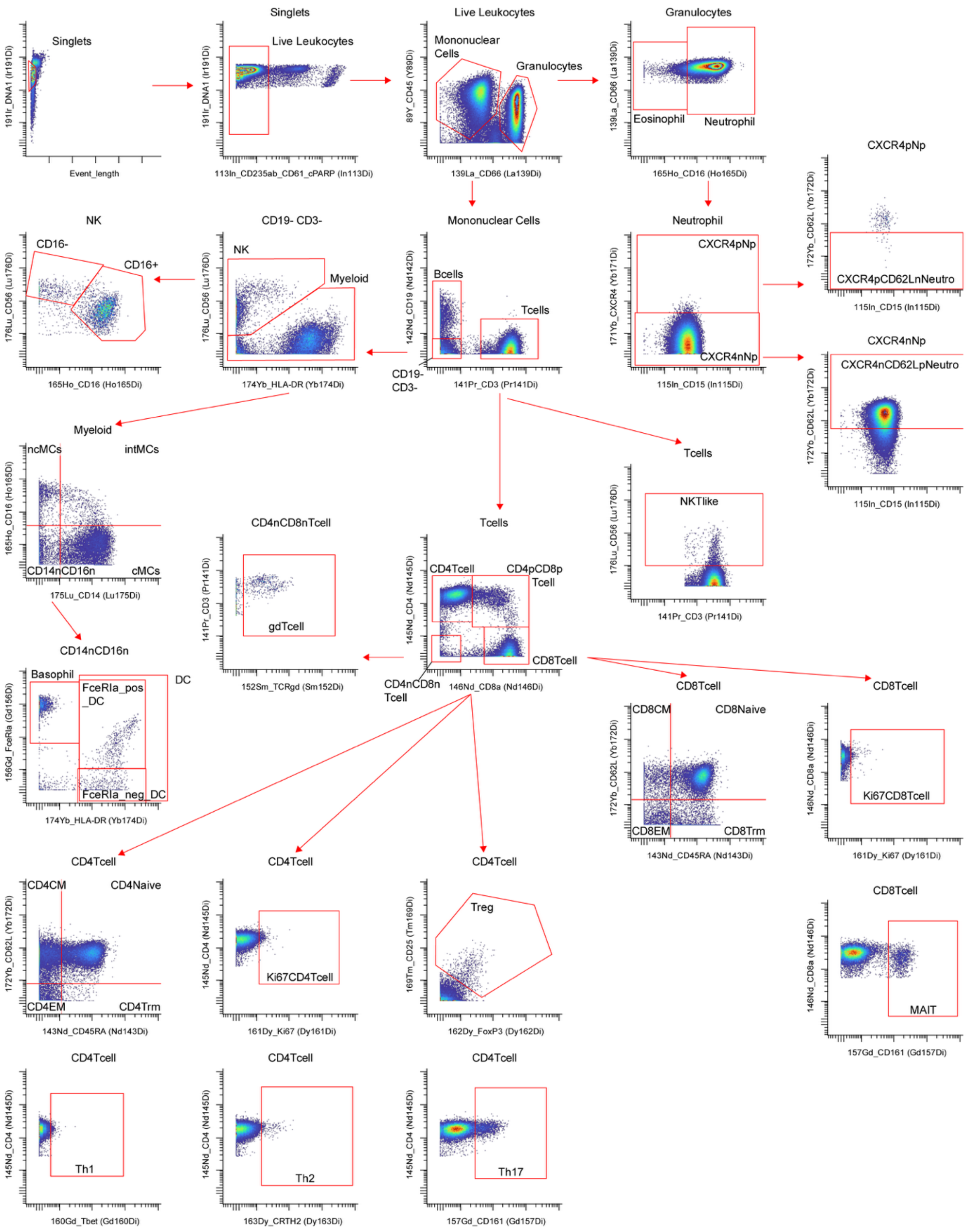


**Figure S1 – Immune cell gating strategy.** Bivariate dot plots for the identification of immune cell populations. Title above each dot plot represents the input gate. NK = natural killer cell; ncMCs = non-classical monocytes; intMCs = intermediate monocytes; cMCs = classical monocytes; DC = dendritic cell; NKTlike = natural killer like T cell; CD8CM/CD4CM = CD8/CD4 central memory T cell; CD8/CD4EM = CD8/CD4 effector memory T cell; CD8/CD4Trm = CD8/CD4 terminal effector memory T cell; MAIT = mucosal-associated invariant T cell; Treg = regulatory T cell

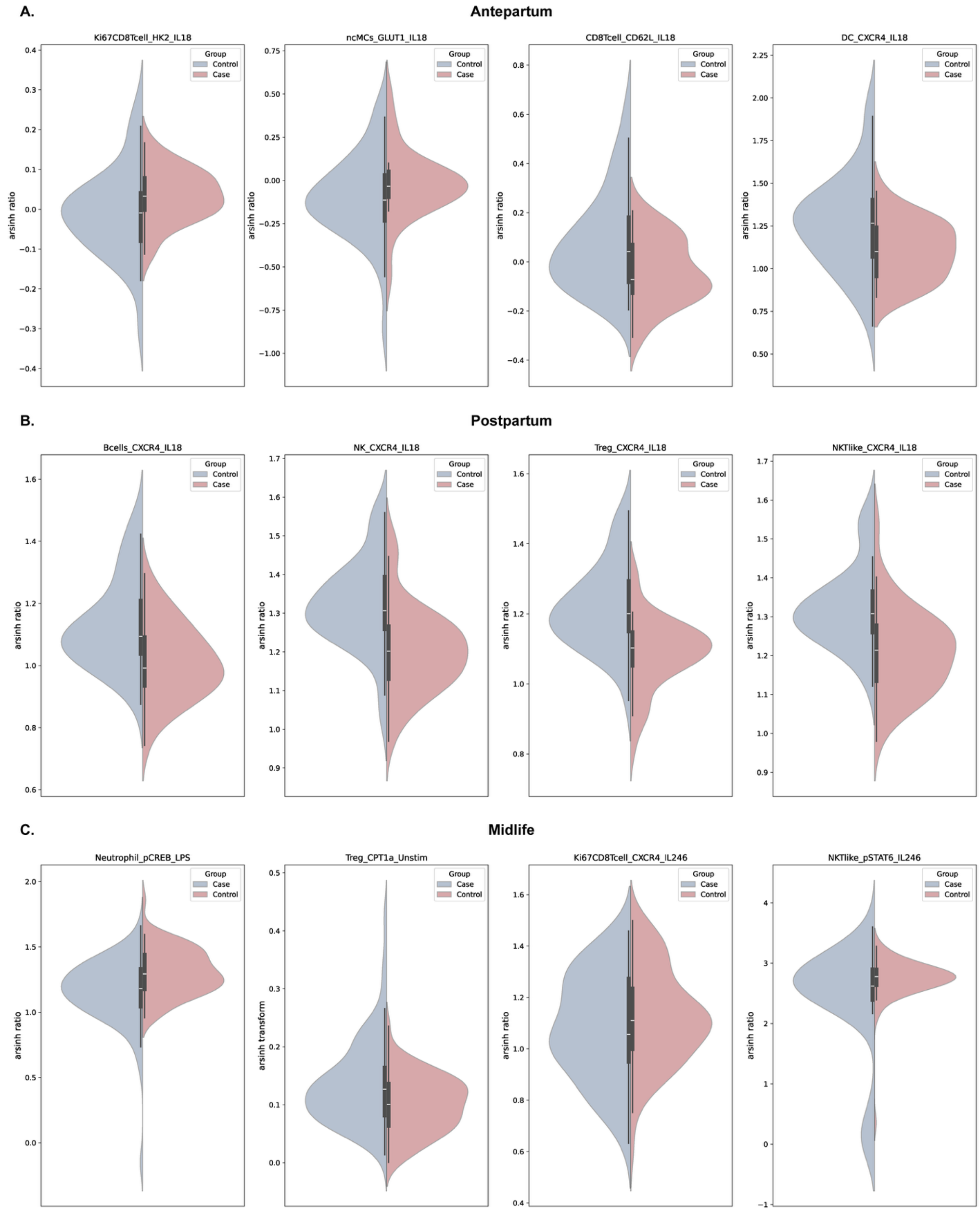


**Figure S2 - Informative model features.**

Violin plots highlight representative model features for each cohort. Values indicate median marker expression. **A.** Selected features in the antepartum model. **B.** Selected features in the postpartum model. **C.** Selected features in the midlife model. ncMCs = non-classical monocytes; DC = dendritic cell; Treg = regulatory T cell.

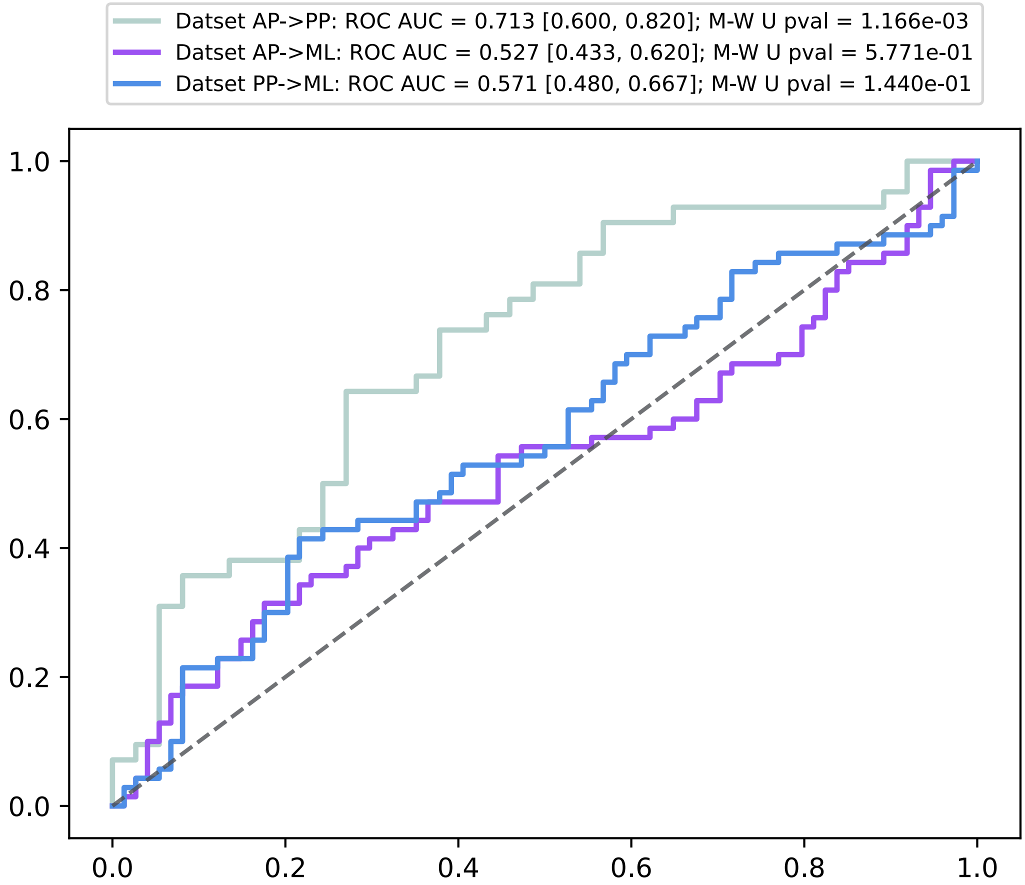


**Figure S3 – Model performance when applied to later timepoints**

*Performance of models when AP or PP models are applied to predict outcomes in later cohorts, displayed as Area Under the Receiver Operator Characteristic curves. AP→PP = AP model used to predict outcomes in PP cohort; AP→ML = AP model used to predict outcomes in ML cohort; PP→ML = PP model used to predict outcomes in ML cohort. AP = antepartum; PP = postpartum; ML= midlife*

**Table S1 - Antibody panel**

| Target | Clone | Vendor | Metal | Concentration used [ug/mL] | Surface/Intracellular staining |
|---|---|---|---|---|---|
| CD45 | HI30 | BioLegend | Y89 | 1 | Surface |
| Barcode 1 | | Trace Sciences | Pd102 | 15μM | Intracellular |
| Barcode 2 | | Trace Sciences | Pd104 | 15μM | Intracellular |
| Barcode 3 | | Trace Sciences | Pd105 | 15μM | Intracellular |
| Barcode 4 | | Trace Sciences | Pd106 | 15μM | Intracellular |
| Barcode 5 | | Trace Sciences | Pd108 | 15μM | Intracellular |
| Barcode 6 | | Trace Sciences | Pd110 | 15μM | Intracellular |
| CD235ab | HIR2 | BioLegend | In113 | 1 | Surface |
| CD61 | VI-PL2 | BD Biosciences | In113 | 0.5 | Surface |
| cPARP | F21-852 | BD Biosciences | In113 | 2 | Intracellular |
| CD15 | W6D3 | BioLegend | In115 | 2 | Surface |
| CD66 | B1.1/CD66 | BD Biosciences | La139 | 1 | Surface |
| total H3 | D1H2 | Cell Signaling Technology | Ce140 | 2 | Intracellular |
| CD3 | UCHT1 | BioLegend | Pr141 | 1 | Surface |
| CD19 | HIB19 | BioLegend | Nd142 | 1 | Surface |
| CD45RA | HI100 | BioLegend | Nd143 | 1 | Surface |
| VDAC1 | 20B12AF2 | Abcam | Nd144 | 2 | Intracellular |
| CD4 | RPA-T4 | BioLegend | Nd145 | 2 | Surface |
| CD8a | RPA-T | BD Biosciences | Nd146 | 1 | Surface |
| GLUT1 | EPR3915 | Abcam | Sm147 | 2 | Surface |
| CPT1a | 8F6AE9 | Abcam | Nd148 | 2 | Surface |
| pCREB | 87G3 | Cell Signaling Technology | Sm149 | 2 | Intracellular |
| pSTAT5 | C11C5 | Cell Signaling Technology | Nd150 | 4 | Intracellular |
| pp38 | 36/p38 (pT180/pY182) | BD Biosciences | Eu151 | 2 | Intracellular |
| TCRγδ | B1.1/CD66 | BD Biosciences | Sm152 | 4 | Surface |
| pSTAT1 | 14/P-STAT1 | BD Biosciences | Eu153 | 1 | Intracellular |
| pSTAT3 | M9C6 | Cell Signaling Technology | Sm154 | 4 | Intracellular |
| pS6 | SP50 | Abcam | Gd155 | 2 | Intracellular |
| FceRIα | AER-37 (CRA-1) | BioLegend | Gd156 | 0.5 | Surface |
| CD161 | HP-3G10 | BioLegend | Gd157 | 4 | Surface |
| PD-L1 | 29E.2A3 | BioLegend | Gd158 | 4 | Surface |
| H3K27me3 | MABI 0323 | Active Motif | Tb159 | 2 | Intracellular |
| Tbet | 4B10 | ThermoFisher | Gd160 | 8 | Intracellular |
| Ki67 | B56 | BD Biosciences | Dy161 | 2 | Intracellular |
| FoxP3 | PCH101 | ThermoFisher | Dy162 | 10 | Intracellular |
| CRTH2 | BM16 | BioLegend | Dy163 | 4 | Surface |
| CD44 | BJ16 | BioLegend | Dy164 | 0.5 | Surface |
| CD16 | 3G8 | BioLegend | Ho165 | 2 | Surface |
| pNFκB | K10-895.12.50 | BD Biosciences | Er166 | 2 | Intracellular |
| pERK1/2 | D13.14.4E | Cell Signaling Technology | Er167 | 2 | Intracellular |
| pSTAT6 | A15137E | BioLegend | Er168 | 2 | Intracellular |
| CD25 | M-A251 | BioLegend | Tm169 | 2 | Surface |
| pPLCg1 | A17025A | BioLegend | Er170 | 4 | Intracellular |
| CXCR4 | 12G5 | BioLegend | Yb171 | 4 | Surface |
| CD62L | DREG.200 | ThermoFisher | Yb172 | 0.25 | Surface |
| HK2 | EPR20839 | Abcam | Yb173 | 4 | Intracellular |
| HLA-DR | L243 | BioLegend | Yb174 | 2 | Surface |
| CD14 | M5E2 | BioLegend | Yb175 | 4 | Surface |
| CD56 | NCAM16.2 | BD Biosciences | Yb176 | 1 | Surface |
| DNA | | Fluidigm | Ir191/193 | 50μM | Intracellular |
| CD63 | H5C6 | BD Biosciences | Bi209 | 0.5 | Surface |

**Table S2 – Confounder analysis**

| | coefficient | Standard error | $t$ value | $Pr > \|t\|$ | Significance |
|---|---|---|---|---|---|
| Antepartum Model | | | | | |
| age | -0.0117277 | 0.00819156 | -1.4316793 | 0.1576026 | NS |
| BMI at study visit | 0.01127746 | 0.00857463 | 1.31521236 | 0.19361458 | NS |
| parity | 0.04357068 | 0.05352212 | 0.81406858 | 0.41893542 | NS |
| study visit timing (GA) | -0.0288742 | 0.01020742 | -2.82875 | 0.00640655 | ** |
| GDM | 0.16399123 | 0.1680492 | 0.97585252 | 0.33319024 | NS |
| systolic | 0.00705574 | 0.00278467 | 2.53377856 | 0.01400802 | * |
| diastolic | 0.0064868 | 0.00568303 | 1.14143336 | 0.25838046 | NS |
| modelPreds | 0.33157814 | 0.13035584 | 2.54363859 | 0.01365808 | * |
| Postpartum Model | | | | | |
| age | -0.0100543 | 0.01322547 | -0.7602203 | 0.4497128 | NS |
| BMI at study visit | 0.00789642 | 0.00976408 | 0.80872132 | 0.42145522 | NS |
| parity | 0.03382038 | 0.06688209 | 0.50567167 | 0.61469881 | NS |
| study visit timing (weeksSinceDelivery) | 0.01221979 | 0.00520066 | 2.3496608 | 0.02165802 | * |
| GDM | 0.41748391 | 0.19546686 | 2.13582956 | 0.03624407 | * |
| systolic | 0.00020336 | 0.0057866 | 0.03514355 | 0.97206674 | NS |
| diastolic | 0.0023361 | 0.00928791 | 0.25152017 | 0.80215957 | NS |
| modelPreds | 0.44871433 | 0.19805219 | 2.26563676 | 0.02661499 | * |
| Midlife Model | | | | | |
| age | -0.0228414 | 0.00756955 | -3.0175404 | 0.00304682 | ** |
| BMI at study visit | 0.00400979 | 0.00839975 | 0.47737095 | 0.63386994 | NS |
| parity | 0.01487917 | 0.03749529 | 0.39682755 | 0.69212125 | NS |
| study visit timing (yearsSinceDelivery) | 0.01491857 | 0.01294151 | 1.15276857 | 0.25104248 | NS |
| GDM | 0.57227264 | 0.19833578 | 2.88537266 | 0.00455278 | ** |
| systolic | 0.00339593 | 0.00472445 | 0.71879884 | 0.47350717 | NS |
| diastolic | 0.00787966 | 0.00610156 | 1.29141571 | 0.19876635 | NS |
| modelPreds | 0.42593933 | 0.16310091 | 2.61150796 | 0.01003427 | * |

Variables examined for potential confounding do not create a major impact on the outcome of the multivariate models. Model predictions remain significant for each cohort when considering potential confounding variables. GA = gestational age; GDM = gestational diabetes mellitus. BMI = body mass index. NS = not significant. Significance codes: * < 0.05, ** < 0.01.

**Table S3 – Informative model features**. Features selected for each model, along with their coefficients in the final model. Negative coefficients indicate decreased values in cases compared to controls.

| AP Model Feature | Coefficient | AP Model Feature - continued | Coefficient |
|---|---|---|---|
| ncMCs_VDAC1_IL18 | 2.408 | FceRIanegDC_CXCR4_IL246 | -0.511 |
| Ki67CD8Tcell_HK2_IL18 | 1.676 | gdTcell_pP38_IL18 | -0.572 |
| ncMCs_GLUT1_IL18 | 1.669 | DC_CXCR4_LPS | -0.597 |
| Treg_HK2_IL18 | 1.524 | Th2_pP38_IL18 | -0.603 |
| Ki67CD4Tcell_H3K27me3_IL18 | 1.268 | DC_VDAC1_Unstim | -0.704 |
| Th2_CXCR4_Unstim | 1.005 | FceRIaposDC_CD62L_LPS | -0.73 |
| Eosinophil_pERK1-2_IL18 | 0.982 | Neutrophil_pP38_IL18 | -0.869 |
| DC_pNFkB_Unstim | 0.637 | CD8Tcell_CD62L_IL18 | -1.574 |
| Neutrophil_GLUT1_LPS | 0.599 | DC_CXCR4_IL18 | -5.101 |
| NKTlike_H3K27me3_IL18 | 0.408 | PP Model Feature | Coefficient |
| Ki67CD8Tcell_CD62L_IL18 | 0.339 | NK_CXCR4_IL18 | 0.035 |
| CD4CM_frequency_Unstim | 0.328 | Bcells_CXCR4_IL18 | -0.659 |
| Treg_HK2_IL246 | 0.322 | NKTlike_CXCR4_IL18 | -0.774 |
| CD8CM_pP38_IL18 | 0.303 | Treg_CXCR4_IL18 | -0.835 |
| Neutrophil_CD62L_LPS | 0.302 | ML Model Feature | Coefficient |
| FceRIaposDC_pERK1-2_LPS | 0.235 | Ki67CD8Tcell_pCREB_IL246 | 1.146 |
| gdTcell_H3K27me3_IL18 | 0.227 | Treg_CPT1a_Unstim | 0.88 |
| DC_VDAC1_LPS | 0.206 | Basophil_pP38_LPS | 0.523 |
| Treg_CPT1a_Unstim | 0.095 | Ki67CD8Tcell_pCREB_IL18 | 0.272 |
| Treg_pPLCg1_Unstim | 0.086 | NKTlike_frequency_Unstim | 0.123 |
| intMCs_GLUT1_IL18 | 0.083 | Granulocytes_CD62L_Unstim | -0.214 |
| CD4Trm_pSTAT1_IL246 | -0.026 | Neutrophil_CD62L_Unstim | -0.346 |
| Bcells_PD-L1_IL18 | -0.064 | Granulocytes_pCREB_LPS | -0.415 |
| Ki67CD8Tcell_CD62L_Unstim | -0.221 | NKTlike_pSTAT6_IL246 | -0.442 |
| CD16+_pCREB_LPS | -0.238 | Neutrophil_pCREB_LPS | -0.477 |
| Th1_CD62L_IL246 | -0.303 | Basophil_pSTAT6_IL246 | -0.545 |
| Neutrophil_VDAC1_Unstim | -0.354 | CD16-_H3K27me3_IL246 | -0.639 |
| DC_pSTAT3_IL246 | -0.483 | Ki67CD8Tcell_CXCR4_IL246 | -0.749 |
